# Changes in cognition and neuroinflammation in a rodent model of chemotherapy-induced cognitive impairment are variable both acutely and chronically

**DOI:** 10.1101/2024.02.22.581658

**Authors:** Olivia J. Haller, Ines Semendric, Lyndsey E. Collins-Praino, Alexandra L. Whittaker, Rebecca P. George

**Author notes:** Corresponding Author Email and Address, School of Biomedicine, The University of Adelaide, North Terrace Campus, Adelaide, South Australia, 5005. Co-first authors (Olivia J. Haller, Ines Semendric).

## Abstract

Chemotherapy-induced cognitive impairment (CICI) affects up to 75% of cancer survivors between 6-months and 20-years post-treatment. Impairments include memory loss, learning difficulties, inability to concentrate and a decrease in processing speed, all of which can negatively impact quality of life. Several mechanisms are proposed to drive these impairments, with evidence implicating neuroinflammation as a key contributor. However, the time course over which impairments occur is less well-established. Several preclinical studies have focused on acute (0-7 days) and sub-acute (8-days to 12-weeks) time-points following chemotherapy, but few have investigated more longer-term time-points. This study therefore aimed to understand the evolution of cognitive changes following methotrexate (MTX) or 5-flurouracil (5-FU) chemotherapy treatment, assessing three time-points: acute (96-hour), sub-acute (31-days) and chronic (93-days). Further, we investigated whether alterations in cognition were associated with concomitant changes in neuroinflammation, assessed via astrocytic reactivity. Female Sprague Dawley rats received two intraperitoneal injections of MTX, 5-FU or saline and were assessed on the novel object recognition, 5-choice serial reaction time task and Barnes maze. Hippocampal (HIPP) and pre-frontal cortex (PFC) tissue was examined for GFAP expression. Results indicate that both MTX and 5-FU exposure were associated with impairments in spatial memory, task acquisition, and processing speed at 31-days, with impairment ameliorated by 93-days post-treatment. While MTX and 5-FU increased GFAP expression across all time-points, with the largest changes at 96-hours and 31-days, drug-specific and region-specific variations were noted. These results provide valuable insight into the complexity of the neuroinflammatory response in CICI. While neuroinflammation may be a promising therapeutic target, further work is required to fully understand this response, including which aspects to target and at what time-points, to ensure optimal outcomes for cancer patients treated with chemotherapy.

## 1. Introduction

The use of chemotherapy for the treatment of cancer has helped reduce recurrence and improve survival rate (1). However, patients that have received chemotherapy often experience debilitating side-effects, both short and long-term (2). In particular, individuals may experience cognitive impairment, a condition termed chemotherapy-induced cognitive impairment (CICI), or, more colloquially, “Chemobrain”. This condition affects a significant proportion of patients, with 1 in 3 breast cancer patients reported to have experienced cognitive impairment post chemotherapy treatment (3). The cognitive domains affected are diverse, with the most common including short-term, working and visual-spatial memory, executive function and attention span (4, 5). These result from structural and functional changes to multiple brain regions, including frontal, prefrontal, parietal and temporal regions (6, 7). It should be noted that, whilst this impairment is often attributed to chemotherapy treatments, there is evidence that cancer alone can contribute to this condition (8, 9). However, animal models assessing the effects in cancer-free rats suggest that chemotherapy itself plays a fundamental role in inducing impairments, regardless of cancer state (10–12). Additionally, cognitive impairment has also been shown to occur in patients receiving chemotherapy as a treatment modality for other conditions, such as rheumatoid arthritis (13). These impairments result in a decreased quality of life for cancer survivors, as they are often unable to return to work, study or maintain a social life, and may struggle to provide and care for their families (14).

Candidate mechanisms identified as drivers of CICI include neuroinflammation (15), impaired neurogenesis (16), oxidative stress (17), apoptosis (18) and blood-brain barrier (BBB) breakdown (19). Of these, neuroinflammation has received considerable focus as a pivotal player in the development of CICI, with animal studies showing an increase in pro-inflammatory cytokines, microglia and astrocytes in various regions of the brain (15, 20). Neuroinflammation is an inflammatory response within the brain or spinal cord. It may result from both direct and indirect stressors, such as chemotherapy exposure, and it, in turn, triggers a cascade of cellular signalling events and consequent neurotoxicity, which can negatively impact cognition via increased neuronal death (for review, see (21)). Inflammation is mediated through the activation of glial cells, such as microglia and astrocytes, and subsequent release of cytokines and other cell signalling molecules (22). While this response acts to repair damage in the acute phase of injury, it can become neurotoxic when it persists long-term (21). Prior studies have demonstrated that CICI patients receiving chemotherapy treatment for cancer may experience such persistent alterations in neuroinflammatory response (20). For example, individuals with CICI have been shown to have increased levels of circulating pro-inflammatory cytokines in serum, such as Il-1B and Il-6 one-month after chemotherapy (23). Similarly, a study investigating paclitaxel chemotherapy administration in C57BL/6 mice found that hippocampal microglia was significantly increased after 5-days of recovery (24). To date, while the role of microglia in CICI has been well-characterised (25), less is known about the function that astrocytes serve in this process. Astrocytes are multifaceted cells that aid in multiple physiological processes related to inflammation, including regulating neuroimmune processes by either perpetuating inflammation or promoting immunosuppression and repair through their ability to become reactive (26). It is thus reasonable to hypothesise that astrocyte reactivity would be up-regulated in CICI. In line with this, the effect of doxorubicin chemotherapy on astrocyte reactivity in male Wistar rats was investigated, and it was concluded that the number, mean intensity and volume of GFAP+ cells were significantly increased compared to control animals in the HIPP (27).

In spite of emerging evidence that neuroinflammation plays a critical role in the brain mechanisms underlying CICI, critical questions remain about the role of reactive astrocytes in this process. Further, the evolution of changes in neuroinflammation over time, and how this relates to the emergence and maintenance of cognitive impairment following chemotherapy exposure, remains unclear. Prior studies in the area have primarily focused on acute time-points (0-7 post chemotherapy treatment) of CICI, with a lack of focus on what is happening at more chronic time-points, leading to a significant gap in the field (28). An understanding of this timeframe is needed in order to develop therapeutic strategies aimed at modulating the neuroinflammatory response, and to determine the optimal time of administration of such therapies. Furthermore, clinically, impairments sometimes do not manifest until several months following treatment, or even beyond (2), making the assessment of long-term time points particularly relevant. Therefore, the aim of this study was to comprehensively characterise the time course of chemotherapy-induced cognitive changes, and to probe concomitant changes in astrocytic reactivity, in a CICI rat model at an acute (96-hour post treatment), sub-acute (31-days post treatment) and chronic time-point (93-days post treatment) following methotrexate (MTX) and 5-fluouracil (5-FU) chemotherapy treatment.

## 2. Methods

### 2.1 Animals

Female Sprague Dawley rats (Hsd: SD; n=108) aged approximately 6-8 weeks, weighing between 150-220g were sourced from a certified Specific Pathogen Free facility (Laboratory Animal Services, The University of Adelaide). Power calculations were made to derive the sample size n=12 per treatment group to achieve 80% power, by using the standard deviation (3.5s) and effect size (0.74) from previous studies that utilised the novel object recognition test (NOR) (29). The rodent populations were free of viral, bacterial, and parasitological infections. SD rats are a commonly used outbred strain and were selected to mimic the heterogeneity existing in the human population (30). Females were selected, as the clinical literature primarily reflects the female breast cancer population, with up to 82% of women thought to experience cognitive impairment from chemotherapy treatment (2). Animals were housed in groups of six in standard polycarbonate open-top cages (415mm x 260mm x 145mm, Tecniplast, Exton, PA, USA). Rats were given cardboard structures and shredded paper as nesting material and enrichment. Acidified RO water and a standard diet of chow (Teklad Irradiated Global SoyProtein-Free Extruded Rodent Diet, Envigo, Madison, WI) was available *ad libitum* throughout acclimation and treatment regimen. Thereafter, food was restricted to 4g/100gbwt throughout behavioural testing for the 31 and 93-day groups to maintain motivation for the sugar pellet reward. Animals were monitored and weighed daily, with additional food given if bodyweight dropped below 15% of pre-feeding weight. Rats were acclimatised to the animal facility for 5-days prior to the commencement of experimental procedures. During this period, animals were handled and given wet food daily. The facility was maintained under a 12-hour light/dark cycle (0700h – 1900h light) and room temperature at 21-23°C.

All experimental procedures were approved by the University of Adelaide Animal Ethics Committee (S-2019-019) and were conducted in accordance with the Australian Code for the Care and Use of Animals for Scientific Purposes, 8^th^ Edition, 2013 guidelines. This paper was written in accordance with the Animal Research: Reporting in vivo experiments: The ARRIVE guidelines (31).

### 2.2 Chemotherapy agents

The agents used were purchased already prepared and included MTX (37.5mg/kg, *Accord Healthcare Pty Ltd*, Melbourne, Australia) and 5-FU (75mg/kg, *Hospira Australia Pty Ltd,* Victoria, Australia), with sterile saline (0.9% Sodium Chloride Inj, *Braun Australia*, Australia) used for the control. Chemotherapy agents were selected based on clinical usage (32, 33) and dosage was based on previous literature that had demonstrated significant cognitive impairment following treatment in animal models (34, 35).

### 2.3 Experimental design and treatment administration

Animals weighing >140g were randomly allocated, using a random number generator, to receive either MTX (37.5mg/kg IP; Accord Healthcare Pty Ltd, Melbourne, Victoria, Australia), 5-FU (75mg/kg IP; Hospira Australia Pty Ltd, Mulgrave, Victoria, Australia) or saline control (equivalent dose to chemotherapy). Animals were then further randomly allocated via an online random number generator to one of three time-points to investigate acute, sub-acute and chronic effects: 96-hours post-treatment (MTX n=12, 5-FU n=12; Saline n=12) (Figure 1A), 31-days post treatment (MTX n=12, 5-FU n=12; Saline n=12) (Figure 1B) or 93-days post treatment (MTX n=12, 5-FU n=12; Saline n=12) (Figure 1C). Rats received a single intraperitoneal (IP) injection of MTX, 5-FU or saline once weekly for two consecutive weeks. Rats in the saline control group received an equivalent volume. Animals were monitored and weighed daily throughout the duration of treatment. To minimise adverse systemic effects of chemotherapy agents and increase nutrient intake, rats were given Nutrigel (High-energy dietary supplement, *Troy Laboratories Pty Ltd*, NSW, Australia) throughout the duration of treatment and for 3-5 days following the last chemotherapy administration. Following behavioural assessment, rats were humanely euthanized via CO2 asphyxiation (individually placed in a CO2 chamber with gradual fill rate of 20% of chamber volume/minute) at one of three terminal end-points (Figure 1). Following humane euthanasia, the hippocampus (HIPP) and prefrontal cortex (PFC) were dissected, snap frozen in liquid nitrogen and stored at −80°C.

**Figure 1.**
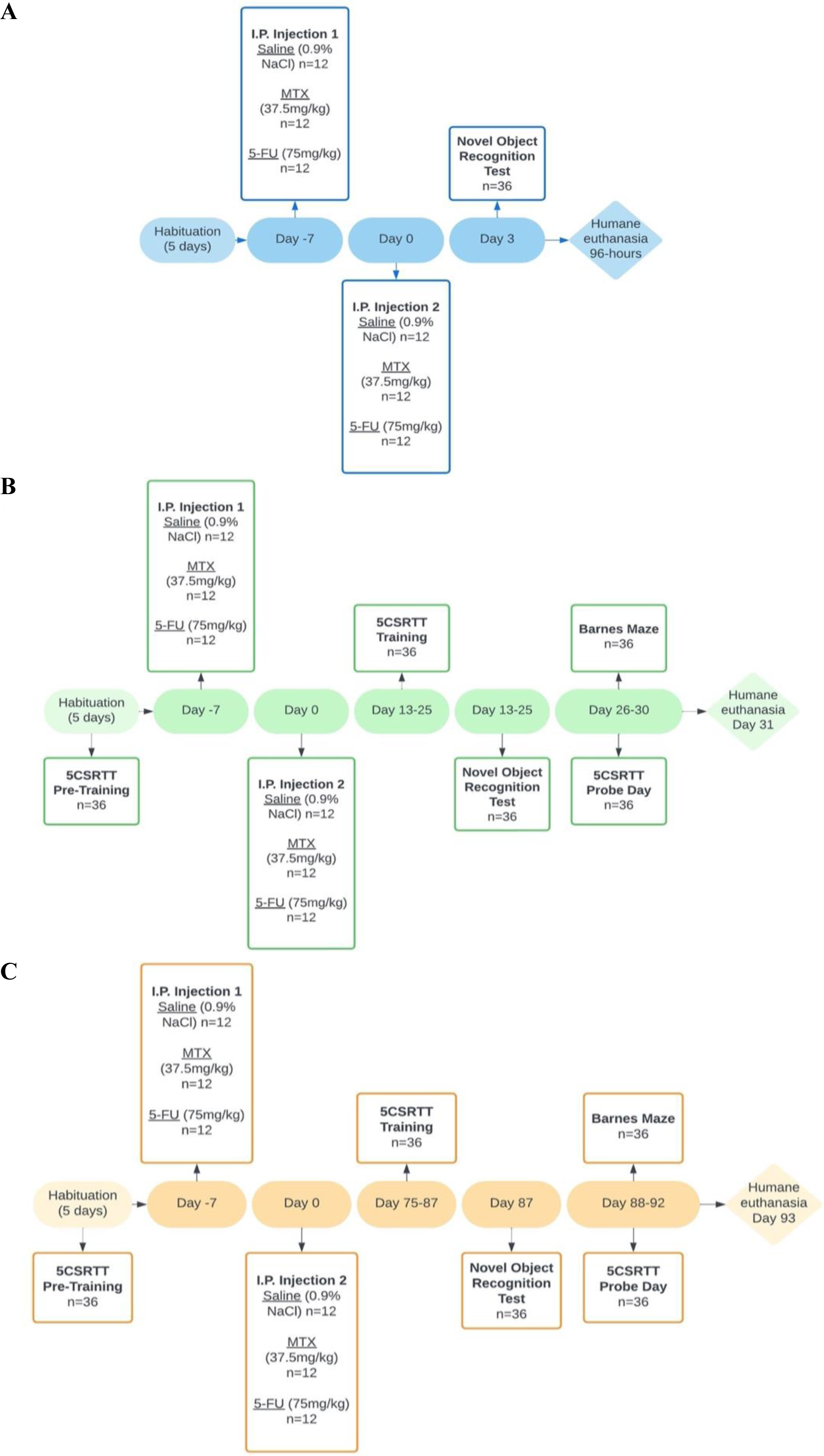
Experimental timeline including chemotherapy administration and behavioural testing at (A) acute time-point (96-hours post treatment), (B) sub-acute time-point (31-days post treatment) and (C) chronic time-point (93-days post treatment) (n=36 per time-point and n=12 per treatment group/time-point).

### 2.4 Cognitive tests

Animals were assessed for cognitive function using the NOR, Barnes maze and five-choice serial reaction time task (5CSRTT) (Figure 1). All animals underwent habituation to behavioural tests. Animals in the sub-acute and chronic time-point groups commenced pre-training for the 5CSRTT prior to commencement of treatment. All behavioural testing was conducted between 0700h – 1900h and was performed in an order of least to most aversive, with appropriate breaks given between behavioural tests. The testing order within each test trial was randomised across animals utilising an online random number generator.

#### 2.4.1 Novel Object Recognition test (NOR)

The NOR was performed at 72 hours, day 25, or day 87 following the last chemotherapy or saline injection (Figure 1). The NOR is a well validated test to assess non-spatial memory and is dependent on the HIPP and perirhinal cortex (36). The ethological basis for this test is that rats with an intact recognition memory will choose to preferentially explore the novel object (36). The testing arena consisted of an enclosed open field (50cm x 50cm x 50cm). All procedures were conducted in a brightly lit room with a video camera (Panasonic Video Camera HC-V180, Selangor, Malaysia) suspended above to record the trials. Animals were firstly habituated to the testing arena without any objects for a 5-minute period. Post habituation, rats were placed into the arena twice, separated by a 1-hour break. During object familiarisation (trial one), two identical objects were placed on equal sides of the arena. The animal was placed in the midpoint of the arena wall opposite from the objects, with their nose facing away from them to prevent any bias in orientation towards one object. The investigator left the room to prevent their presence being used as an external cue. The animal was allowed to freely explore for 5-minutes, before being removed and returned to the home cage for 1-hour. Time spent interacting with each object and total exploration time were recorded in trial one. The object recognition phase (trial two) was conducted identically to trial one; however, one of the objects was replaced with a novel object. The novel object was distinctive from the original object (e.g. colour and shape). The arena and objects were cleaned with 70% ethanol between tests to avoid odour cues. Time spent interacting with the novel object, the familiar object, and total exploration time were recorded in trial two. Retrospective video analysis was conducted using ANY-maze™ software (Stoeltingco, Wood Dale, IL). A preference index was calculated to determine an animal’s preference to the novel versus the familiar object. The preference index for the novel object was determined by a blinded observer utilising the following formula: PI = (time spent with the novel object/time spent with both novel and familiar objects) x 100.

#### 2.4.2 Barnes maze

The Barnes maze is a hippocampal dependent task used to test spatial learning and memory (37) and was performed over five consecutive days, on days 26-30 or 88-92 (Figure 1). The apparatus consisted of a large round platform (1.2m diameter) with eighteen holes (5cm diameter) equally spaced around the circumference of the maze. An escape box was located beneath one of the holes, whilst the remaining holes were false bottomed. All trials were performed in a dimly lit room, with a bright light used to illuminate the surface of the maze to act as a mild negative stimulus to motivate the animal to find the escape box. During the training phase, all animals underwent two trials per day, with a 15-minute break between trials, over a 4-day period. The animal was placed in the centre of the maze and free to explore the apparatus. If the rat located the escape box during the test timeframe, the light was turned off and it was subsequently returned to the home cage. If the rat did not enter the box after 3 minutes, it was gently guided to the escape box by the experimenter, the light was immediately turned off, and the rat was left in the escape box for 20-seconds prior to being returned to the home cage. This was repeated over the subsequent training days. On day 5 (probe trial), the location of the escape box was rotated by 90° from the original position. The trial was conducted in the same manner as the training trial, with a 30-minute break between probe trials. Location of the escape hole was randomised and counterbalanced across treatment groups and time-points. The maze and escape box were cleaned with 70% ethanol between each trial. All trials were video recorded and retrospectively analysed by a blinded observer. Latency to find the escape box during training trials, latency to old and new location of the escape box and number of revisits to the old escape box location during both probe trials were recorded.

#### 2.4.3 Five-Choice Serial Reaction Time Task (5CSRTT)

The 5CSRTT is a hippocampal and PFC dependent task designed to evaluate executive function, which includes attention, inhibition and impulsivity in rodents (38). The 5CSRTT was performed in the Bussey-Saksida Touchscreen operant chamber (Campden Instruments Ltd., England) as previously described (39, 40). Sugar pellets (Dustless Precision Pellet-Sugar Formulation 45mg, ASF0042, Able Scientific, Australia) were utilised as a reward and were dispensed when the animal selected the correct response. The touchscreen operant system was linked to a computer using the ‘Whisker Server’ software to simultaneously operate and control the chambers. ABETII software with pre-set programs was used to measure and record training sessions and probe trials during the 5CSRTT task in each chamber. The ABETII software automatically controlled outcome measurements, presentation of the stimulus, house light, tone generation and dispensing of the sugar pellets.

The 5CSRTT was performed over four phases, which included habituation, pre-training, training, and probe trials (Figure 2). Each animal received one 5CSRTT session per day. Prior to the commencement of chemotherapy and saline treatment, rats were habituated to the testing apparatus and reward system and underwent pre-training for 5-days. During habituation, animals were placed in the operant chambers with 10 sugar pellets in the dispenser for 30-minutes. During pre-training, animals underwent ‘initial touch’ and ‘must touch’ training, where they were trained to nose poke the correct touchscreen in response to the stimulus presented. Rats were required to complete 20 trials in 30-minutes to pass criterion for ‘initial touch’ and 20 trials in 30-minutes for two consecutive days to pass criterion for ‘must touch’.

**Figure 2.**
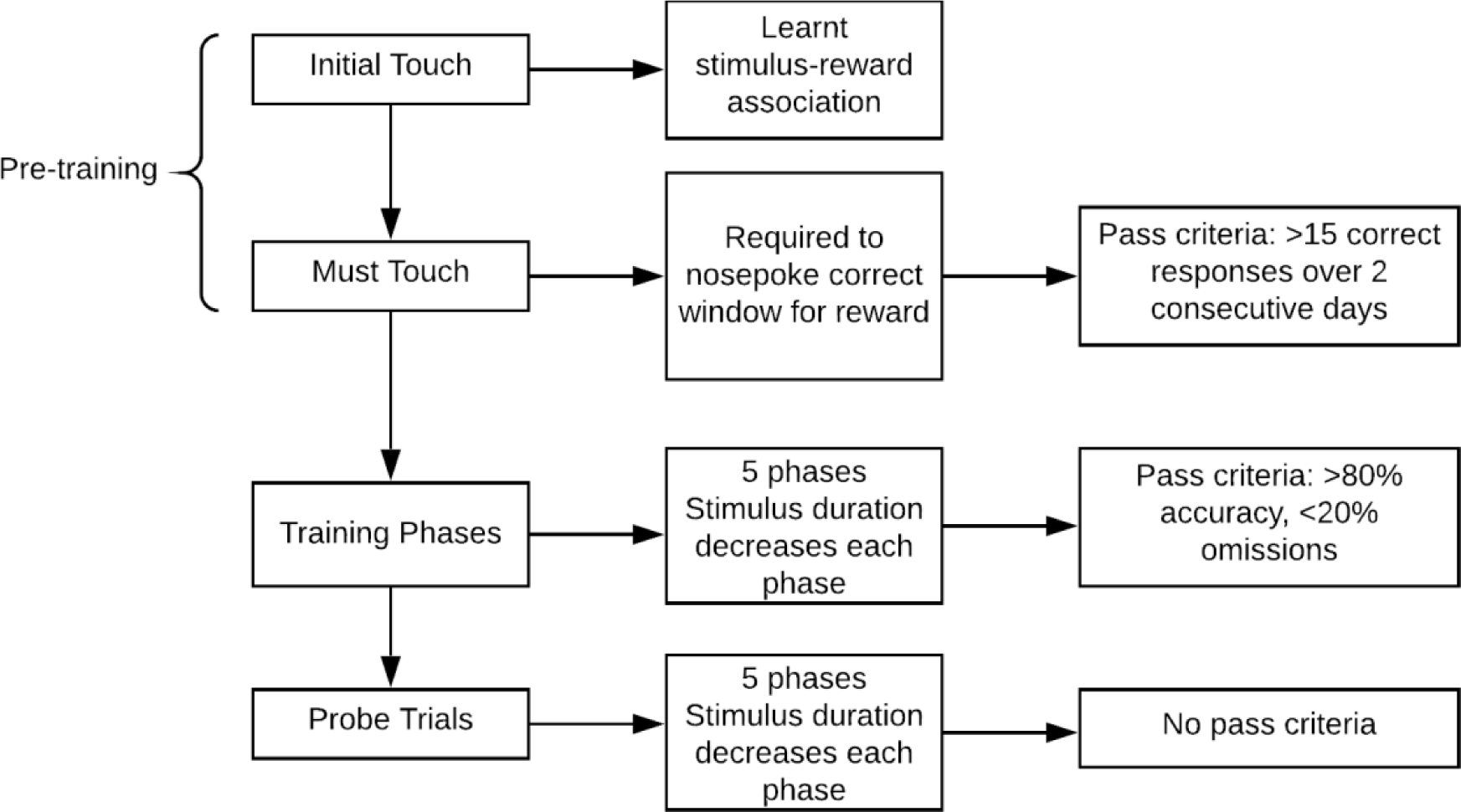
Five-Choice Serial Reaction Time Task (5CSRTT) testing procedure.

On completion of pre-training, animals were moved to the training phase, where they were taught to respond to the correct stimulus in decreasing stimulus durations (60, 30, 20, 10 and 5-seconds) over 13-days. During each session, a sugar pellet was released, and the food dispenser switched on. Once the animal retrieved the pellet, the dispenser light switched off. Then a stimulus was presented within any one of the 5 windows. The animal was required to nose poke the correct window within the specified time. If the right window was poked, a tone was generated, a sugar pellet was released and the food dispenser light turned on. Each session finished at 100 trials or when 30-minutes had elapsed, whichever occurred first. Criteria to pass each training session required rats to achieve at least 80% accuracy and less than 20% omissions. Animals that achieved all training criteria early received retention training until probe trials.

Upon completion of training, animals underwent 5-days of probe trials, which did not have a specific pass criterion (Table 1). During this phase, the stimulus duration was reduced between each trial (4, 2.5, 1.5, 1, 0.5 seconds). The number of trials, accuracy, omissions, premature responses, and correct response latency was analysed.

**Table 1.** Definitions of 5CSRTT outcomes.

| 5CSRTT Variables | Definition |
| --- | --- |
| Number of trials | One trial is defined as the presentation of a stimulus and the animal responding/not responding in the allocated stimulus duration |
| Accuracy | Number of correct responses divided by the sum of correct and incorrect responses, expressed as a percentage (%) |
| Omissions | The total number of no responses to a stimulus, divided by the number of total trials, expressed as a percentage (%) |
| Premature Responses | The total number of responses that occur before the stimulus is presented, divided by the total number of trials |
| Correct Response Latency | The average time from onset of the stimulus to correct nose poke |

### 2.5 Western blot analysis

Levels of glial fibrillary acidic protein (GFAP), a marker of astrocytic reactivity, were measured in the HIPP and PFC. Each tissue sample was cut into 1-2mm sections and homogenised in standard RIPA buffer (Tris-base saline, pH 7.5-8; 50mM, 5M NaCl 150mM, 1% NP-40, 0.5% 10% sodium deoxycholate, 0.1% 10% SDS) with a protease inhibitor tablet (cOmplete^TM^, Mini, EDTA-free Protease Inhibitor Cocktail) using a handheld homogeniser. Samples were sonicated for 3×10-second bursts, followed by 1-minute on ice, then centrifuged at 15,000rpm for 20 minutes at 4°C. The protein concentration was determined using a BCA protein assay kit (Pierce® BCA Protein Assay Kit; 23225, Thermo Fisher Scientific Inc., Victoria, Australia). Gel electrophoresis was performed using Bolt 4–12% Bis-Tris Plus gels (Life Technologies) with 25μg of protein loaded per well. Gels were run at 150V for approximately 1.5-hours or until the dye reached the bottom of the gel. Gels were transferred to a polyvinylidene fluoride (PVDF) membrane using the iBlot 2 Dry Blotting System (Thermo Fisher Scientific Inc). Membranes were stained with Ponceau S red solution (Fluka Analytical) to allow for visual inspection and then washed three times on an agitator for 5-minutes each using 1X tris-buffered saline with tween (TBST). Membranes were then blocked in 5% skim milk solution (5g skim milk powder, 100mL TBST) for 2-hours at room temperature. Following this, membranes were washed three times and incubated at 4°C overnight in 2% skim milk solution (1g skim milk powder, 50mL TBST) with the primary antibody (GFAP; #ab7260, Abcam, Cambridge, UK) and housekeeper protein (GAPDH; #ab8245 Abcam, Cambridge, UK) on a rotating device. The following day, membranes were washed three times in TBST. Membranes were then incubated with secondary antibodies (LICOR 800CW donkey/anti-rabbit,1:10,000; LICOR 800CW donkey/anti-mouse, 1:10,000) in 2% skim milk solution for 2-hours at room temperature on a rotating device covered with aluminium foil. Western blot visualisation was conducted using the Odyssey Infrared Imaging System (model 9120; software version 3.0.21) (LI-COR, Inc.) and analysis was performed by a blinded observer using ImageJ software (Wayne Rashband, National Institutes of Health, USA), where bands were normalised to the housekeeper gene and presented as GFAP relative density.

### 2.6 Statistical analyses

Data were analysed using IBM SPSS (SPSS Inc., Chicago, IL, USA) and Megastat Excel Add-In (version 10.3 Release 3.1.6 Mac, McGraw-Hill Higher Education, New York, NY, USA) statistics software. Four animals were excluded from the study due to humane endpoint implementation, rendering the following sample sizes per treatment group: 96-hour: Saline (n=12), MTX (n=12), 5-FU (n=12); 31-days: Saline (n=11), MTX (n=11), 5-FU (n=11); 93-days: Saline (n=12), MTX (n=11), 5-FU (n=12). Data were checked for normality using the Shapiro–Wilk test. NOR and Western Blot data were analysed using a one-way analysis of variance (ANOVA) with Tukey’s *post-hoc* test. The Barnes maze and 5CSRTT data between groups at each time-point were analysed non-parametrically using a Kruskal-Wallis U-test with *post-hoc* Mann Whitney U-test for task acquisition, latency to old and new escape box on probe day, and the number of sessions required to pass criteria during training phase of the 5CSRTT. Within group differences across time were analysed using a Friedman test with *post-hoc* Wilcoxon signed-rank test. A two-way repeated measures ANOVA with treatment as a between-subjects factor and time as a within-subjects factor with a Bonferroni correction was performed for outcomes assessed on probe day of the 5CSRTT. Data are presented as mean ± SEM for parametric data and median with interquartile range for non-parametric data. A p<0.05 was regarded as statistically significant.

## 3. Results

### 3.1 Effect of MTX and 5-FU treatment on object recognition memory in the NOR

Object recognition memory was assessed utilising the NOR. There was no significant effect of treatment on object recognition memory, as assessed by the preference index for the novel object during trial two, at 96-hours (F(2,27) = 0.46, p=0.6358), 31-days (F(2,29) = 0.412, p=0.67) or 93-days post treatment (F(2,29) = 1.246, p=0.30).

### 3.2 Effects of MTX and 5-FU treatment on memory and learning in the Barnes maze test

During the learning acquisition phase of the Barnes maze test, animals demonstrated no significant difference between treatment groups in daily average escape latency in the sub-acute cohort for Day 1 (Trial 1 (H(2) = 0.291, *p=*0.865), Trial 2 (H(2) = 1.861, *p=*0.394)), Day 2 (Trial 1 (H(2) = 0.022, *p=*0.989), Trial 2 (H(2) = 4.780, *p=*0.092)), Day 3 (Trial 1 (H(2) = 0.920, *p=*0.631), Trial 2 (H(2) = 5.600, *p=*0.061)) or Day 4 (Trial 1 (H(2) = 1.870, *p=* 0.393), Trial 2 (H(2) = 0.088), *p=*0.957)) (Figure 3A). However, a Friedman analysis revealed a significant difference across time for latency to the escape box during task acquisition both at the sub-acute (Saline (χ^2^ (2) = 18.597, p=0.010); MTX (χ2(2) = 27.154, p = <0.01); 5-FU (χ2(2) = 44.528, p = <0.001)) and the chronic (Saline (χ^2^ (2) = 26.099, p<0.001); MTX (χ2(2) = 46.697, P = <.001); 5-FU (χ2(2) = 22.546, p = 0.002)) time-points for all three groups. Specifically, at the sub-acute time-point, there was a significant improvement in daily average escape latency from acquisition Day 1 Trial 1 compared to Day 1 Trial 2 for rats in all treatment groups (SAL *Z* = −2.805, p = 0.005; MTX *Z* = −1.988, p = 0.047; 5-FU *Z* = −2.934, p = 0.003). No significant differences were found between trials on Day 2, 3, or 4 (p = >0.05).

**Figure 3.**
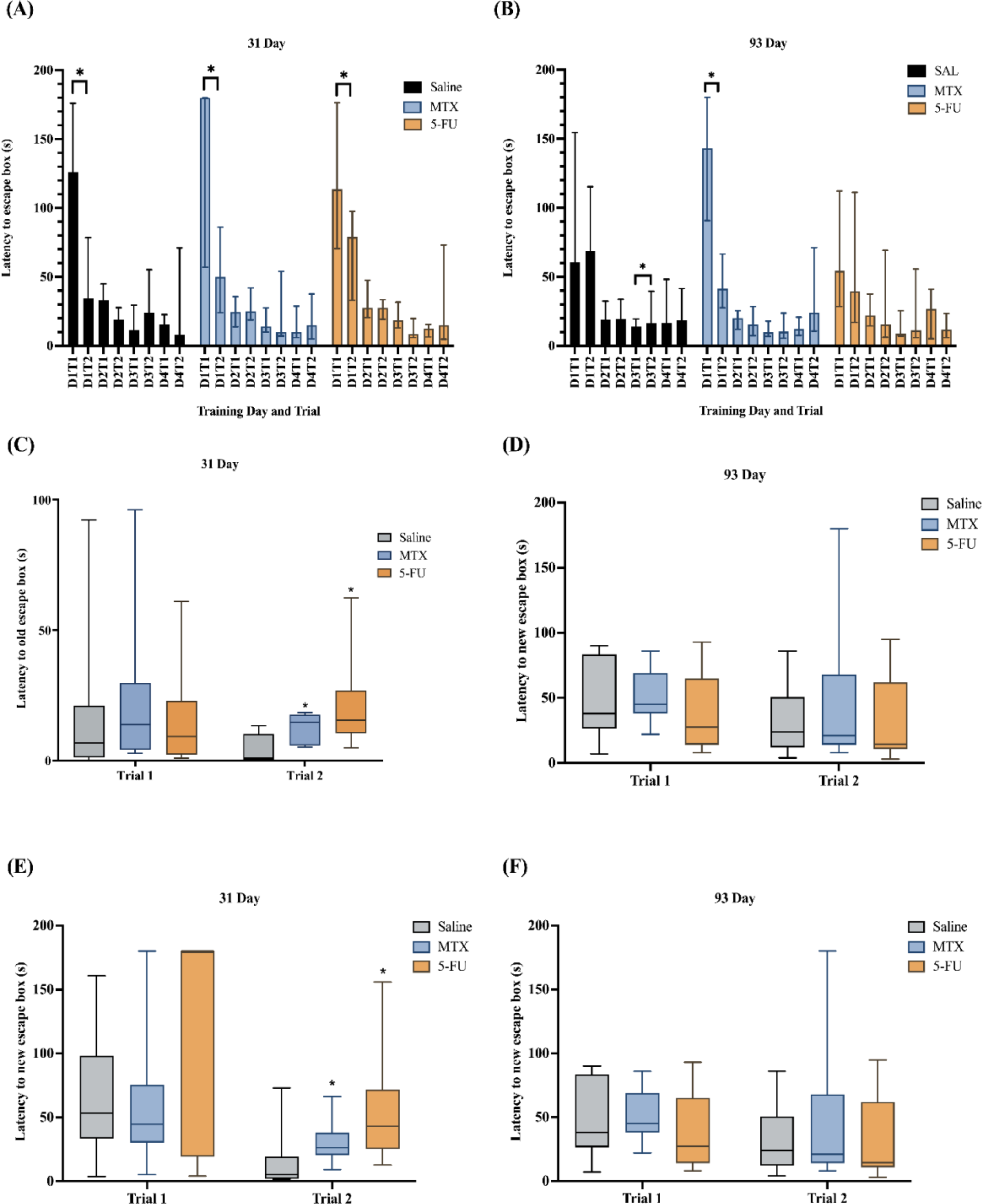
Latency to escape box on acquisition days 1-4 (training) at (A) 31-days and (B) 93-days post-treatment. All animals significantly improved between Day 1 Trial 1 and Day 1 Trial 2 training at 31-days, but only MTX-treated animals improved between Day 1 Trial 1 and Day 1 Trial 2 at 93-days, with SAL animals also improving between Day 3 Trial 1 and Day 3 Trial 2. Latency to old escape box at (C) 31-days was significantly higher for animals treated with MTX and 5-FU, whilst no differences were observed at (D) 93-days post-treatment. Latency to new escape box at (E) 31-days was significantly higher for animals treated with MTX and 5-FU, whilst no differences were observed at (F) 93-days post-treatment on probe day. Data is presented as median with interquartile range. 31-days: Saline (n=11), MTX (n=11), 5-FU (n=11); 93-days: Saline (n=12), MTX (n=11), 5-FU (n=12). * p<0.05 compared to saline within the same trial. # p < 0.05 statistical comparison within group across time.

At 93-days, only MTX treated animals significantly improved between Day 1 Trial 1 and Day 1 Trial 2 (SAL *Z =* −.314, p = 0.754; MTX *Z =* −2.934, p = 0.003; 5-FU *Z =* −.314, p = 0.754). Interestingly, saline-treated animals also improved between Day 3 Trial 1 and Day 3 Trial 2, while neither MTX nor 5-FU groups showed the same pattern (SAL *Z =* −2.318, p = 0.020; MTX *Z =* −.847, p = 0.397; 5-FU *Z =* −1.070, p = 0.875) (Figure 3B). No significant differences were found between trials on Day 2 or Day 4 (p = >0.05).

During the probe trial at the sub-acute time-point, no significant differences were observed between treatment groups on latency to find either the old escape box or the new escape box for probe trial 1 (H (2) = 1.085, p = 0.581; H (2) = 1.356, p=0.508, respectively) (Figure 3C and 3E, respectively). However, on trial 2 on probe day, there was a significant effect of treatment on latency to find the location of the old box (H (2) = 6.155, p=0.046) (Figure 3C). Animals treated with either MTX (mean = 12.32s, SD = 6.11s) or 5-FU (mean = 21.40s, SD = 16.54s) took significantly longer to find the old escape box location compared to saline controls (mean = 3.8s, SD = 6.35s) (MTX vs. SAL: p=0.050; 5-FU vs. SAL: p=0.026; Figure 3C). Likewise, there was also a significant effect of treatment on latency to find the location of the new escape box on trial 2 on probe day (H (2) = 10.947, p=0.004) (Figure 3E). Compared to saline treated animals, animals treated with either MTX (mean = 30.48s, SD = 16.09s) or 5-FU (mean = 50.45s, SD = 40.16s) took significantly longer to find the new escape box location than saline-treated controls (mean = 15.02s, SD = 20.91s) (MTX vs, SAL: p=0.011; 5-FU vs SAL: p=0.004; (Figure 3E). Interestingly, at the chronic time-point, no significant treatment effects were found on latency to find either the old or new escape box location during either the first (F(2,19 = 0.025, p=0.976), (F(2,32 = 1.154, p=0.328), respectively) or second probe trial (F(2,9 = .415, p=0.672), (F(2,32 = .485, p=0.620), respectively) (Figure 3D and 3F). There was also no significant effect between treatment groups for the number of preservative errors made (i.e. number of revisits to the old escape box) during probe trials at either the sub-acute (H (2) = 1.221, p=0.543) or chronic time-points (H (2) = 0.257, p=0.879) (data not graphically represented).

### 3.3 Effect of MTX and 5-FU treatment on performance in the 5CSRTT

Learning was assessed during the training phase of the 5CSRTT. A significant treatment difference was observed in the number of sessions required to pass training criteria at the sub-acute time-point (H (4) = 136.127, *p*<0.001). Post-hoc Mann Whitney U-test revealed that animals treated with 5-FU required a significantly greater number of sessions to reach the 20s stimulus duration (*U =* 31, *p*=0.019), 10s stimulus duration (*U =* 26, *p*=0.018) and 5s stimulus duration (*U =* 25.5, *p*=0.020), compared to saline controls. 5-FU treated animals also took significantly longer to reach the 10s stimulus duration compared to MTX-treated (*U =* 27, *p*=0.041) (Figure 4A). No treatment differences were noted at the chronic time-point for the number of sessions required to pass training criteria for the 60s stimulus duration (H(2) = 0.024, *p=*0.988), 30s stimulus duration (H(2) = 0.218, *p=*0.897), 20s stimulus duration (H(2) = 0.192, *p=*0.909), 10s stimulus duration (H(2) = 3.067, *p=*0.216) or 5s stimulus duration (H(2) = 2.400, *p=*0.301) (Figure 4B).

**Figure 4.**
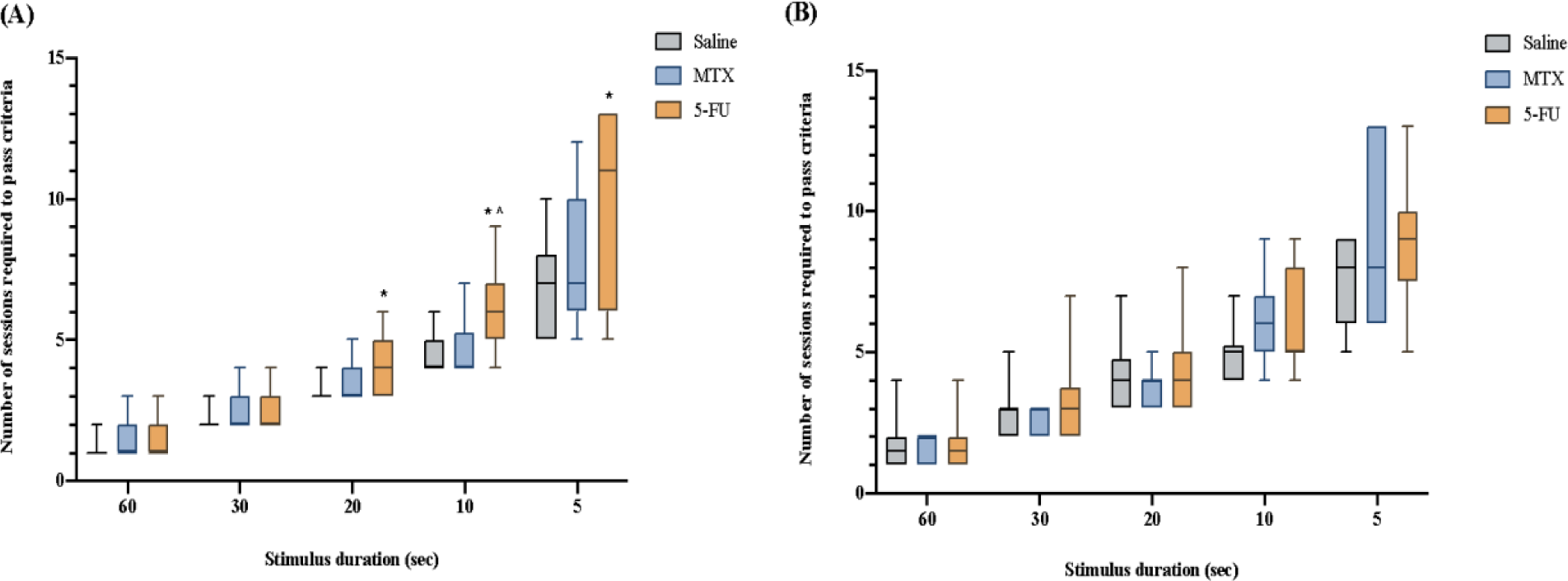
Sessions required to pass criteria during training session at (A) 31-days and (B) 93-days post treatment. Animals treated with 5-FU (n=11) at 31-days required significantly more sessions to pass the trial criteria as the stimulus duration decreased (20s, 10s, 5s). Data represented as median with interquartile range. * p<0.05 compared with saline, × p<0.05 compared with MTX. 31-days post treatment: Saline (n=11), MTX (n=11), 5-FU (n=11); 93-days post treatment: Saline (n=12), MTX (n=11), 5-FU (n=12).

During probe testing, the number of trials completed, accuracy, omissions, premature responses and correct response latency were analysed (Table 1). No significant treatment differences between groups were found at 31-days for accuracy (4s stimulus (H(2) = 1.705, p=0.426); 2.5s stimulus (H(2) = 0.671, p=0.715); 1.5s stimulus (H(2) = 0.473, p=0.789); 1s stimulus (H(2) = 1.203, p=0.548); 0.5s stimulus (H(2) = 1.093, p=0.579)) (Figure 5A), omissions (4s stimulus (H(2) = 2.616, p=0.270); 2.5s stimulus (H(2) = 5.055, p=0.080); 1.5s stimulus (H(2) = 0.576, p=0.750); 1s stimulus (H(2) = 0.483, p=0.785); 0.5s stimulus (H(2) = 2.565, p=0.277)) (Figure 5C), premature responses (4s stimulus (H(2) = 2.493, p=0.288); 2.5s stimulus (H(2) = 1.221, p=0.543); 1.5s stimulus (H(2) = 0.107, p=0.948); 1s stimulus (H(2) = 0.686, p=0.710); 0.5s stimulus (H(2) = 3.646, p=0.162)) (Figure 5E) or correct response latency for most stimulus durations (4s stimulus (H(2) = 1.412, p=0.494); 2.5s stimulus (H(2) = 0.696, p=0.706); 1.5s stimulus (H(2) = 1.110, p=0.574); 0.5s stimulus (H(2) =0.685, p=0.710)) (Figure 5G). However, for correct response latency, there was significance at the 1s stimulus duration (H(2) = 6.540, p=0.038), with 5-FU treated animals demonstrating slower decision-making speed compared to MTX treated animals (MTX vs 5-FU U = 10, p=0.006) (Figure 5G).

**Figure 5.**
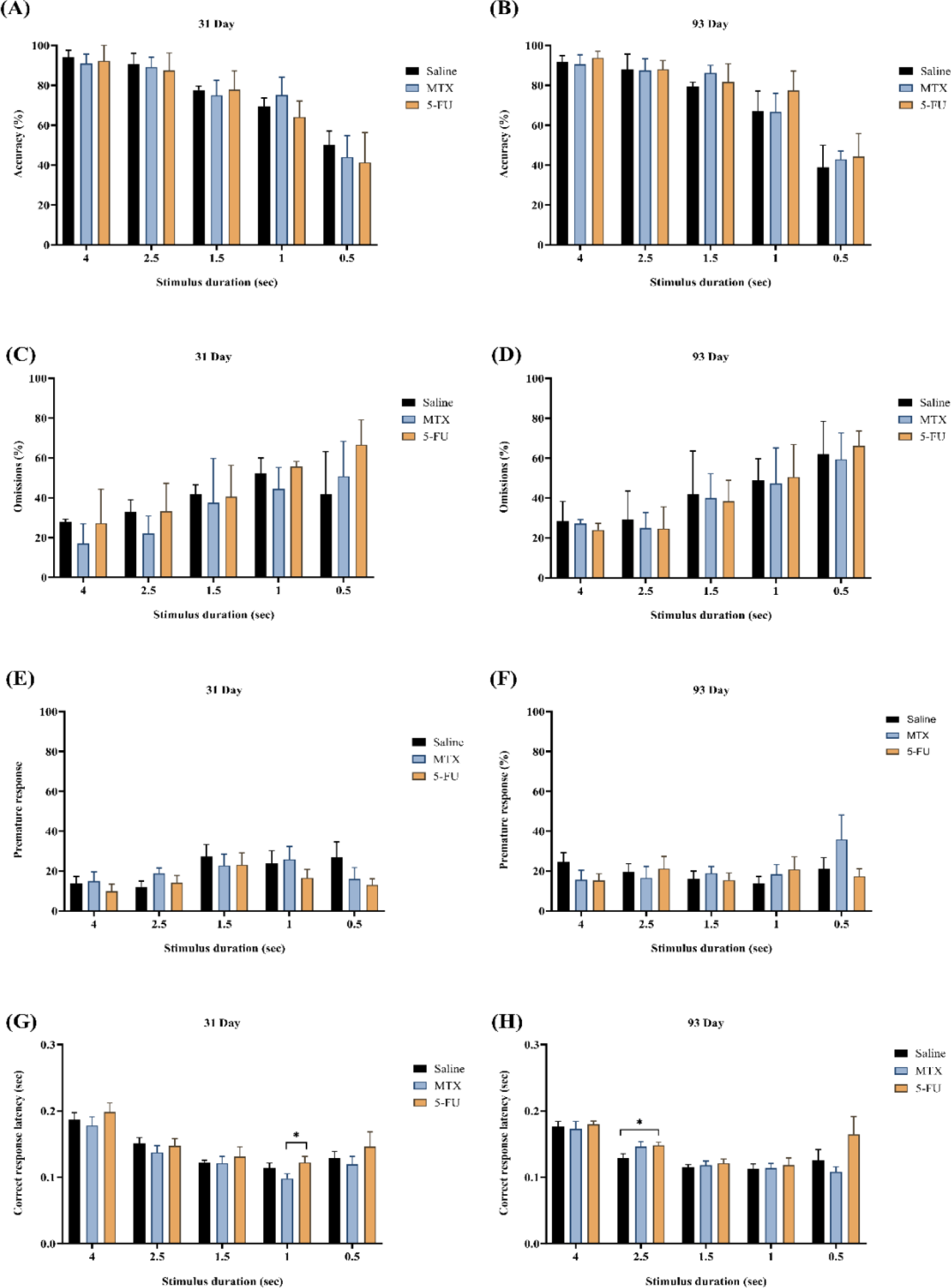
There were no significant effects of chemotherapy and saline on performance in the 5-choice serial reaction time task as assessed by accuracy (A-B), omissions (C-D), premature response (E-F) and correct response latency (G-H), at either 31-days or 93-days post treatment, except between MTX and 5-FU for Correct Response Latency at 1s stimulus duration at 31-days, and between SAL and MTX for Correct Response Latency at 2.5s stimulus duration at 93-days. Data presented as mean ± SEM. 31-days post treatment: Saline (n=11), MTX (n=11), 5-FU (n=11); 93-days post treatment: Saline (n=12), MTX (n=11), 5-FU (n=12). Neither MTX nor 5-FU chemotherapy caused a functionally significant impairment in any of the variables analysed.

Similarly, no significant differences were noted at 93-days for accuracy (4s stimulus (H(2) = 2.705, p=0.259); 2.5s stimulus (H(2) = 0.473, p=0.789); 1.5s stimulus (H(2) = 4.964, p=0.084), 1s stimulus (H(2) = 4.301, p=0.116); 0.5s stimulus (H(2) = 0.509, p=0.775)) (Figure 5B), omissions (4s stimulus (H(2) = 1.382, p=0.501); 2.5s stimulus (H(2) = 0.947, p=0.623); 1.5s stimulus (H(2) = 0.238, p=0.888); 1s stimulus (H(2) = 0.087, p=0.957); 0.5s stimulus (H(2) = 0.115, p=0.944)) (Figure 5D), premature responses (4s stimulus (H(2) = 4.301, p=0.116); 2.5s stimulus (H(2) = 1.305, p=0.521); 1.5s stimulus (H(2) = 1.058, p=0.589); 1s stimulus (H(2) = 0.293, p=0.864; 0.5s stimulus (H(2) = 0.445, p=0.800)) (Figure 5F) or correct response latency for the majority of stimulus durations (4s stimulus (H(2) = 0.904, p=0.636); 1.5s stimulus (H(2) = 0.246, p=0.884); 1s stimulus (H(2) = 0.081, p=0.960); 0.5s stimulus (H(2) = 1.476, p=0.478)) (Figure 5H). However, for correct response latency, there was significance at the 2.5s stimulus duration H(2) = 6.219, p=0.045), with 5-FU treated animals demonstrating slower decision-making speed compared to saline treated animals (SAL vs 5-FU, U = 34, p=0.028) (Figure 5H).

### 3.4 Effect of MTX and 5-FU on astrocytic reactivity marker expression changes in the hippocampus and prefrontal cortex

GFAP expression was used to assess astrocytic reactivity marker expression changes in the HIPP and PFC at 96-hours, 31-days and 93-days post-treatment (Figure 6). Within the HIPP, western blot analysis demonstrated a significant treatment effect in the expression of GFAP at 96-hours (F(2,14 = 7.321, p=0.007). Post-hoc analysis determined that there was a significant increase in both the 5-FU treated group compared to saline controls (p=0.015) and in the MTX treated group compared to saline controls (p=0.013). No significant difference was observed between 5-FU and MTX treated animals (p>0.977) (Figure 6A). At 31-days, there was also a significant treatment effect found (F(2,14 = 14.783, P=< 0.001). There was a significant increase in the 5-FU treated group compared to saline controls (p< 0.001) and in the MTX treated group compared to saline controls (p=0.023). Although no statistically significant difference was observed between 5-FU and MTX treated animals (p=0.059), there was a trend for GFAP levels to be higher in 5-FU treated animals than in MTX-treated animals (Figure 6B). Finally, at 93-days, there was a significant treatment effect found (F(2,14 = 7.134, p=0.007), with GFAP levels higher in the 5-FU treated animals than in those who were administered saline (p=0.006). While there was also a trend for levels to be higher in 5-FU treated animals compared to those treated with MTX, this did not reach statistical significance (p=0.058). Further, MTX-treated animals were not significantly different than saline controls (p=0.444) (Figure 6C).

**Figure 6.**
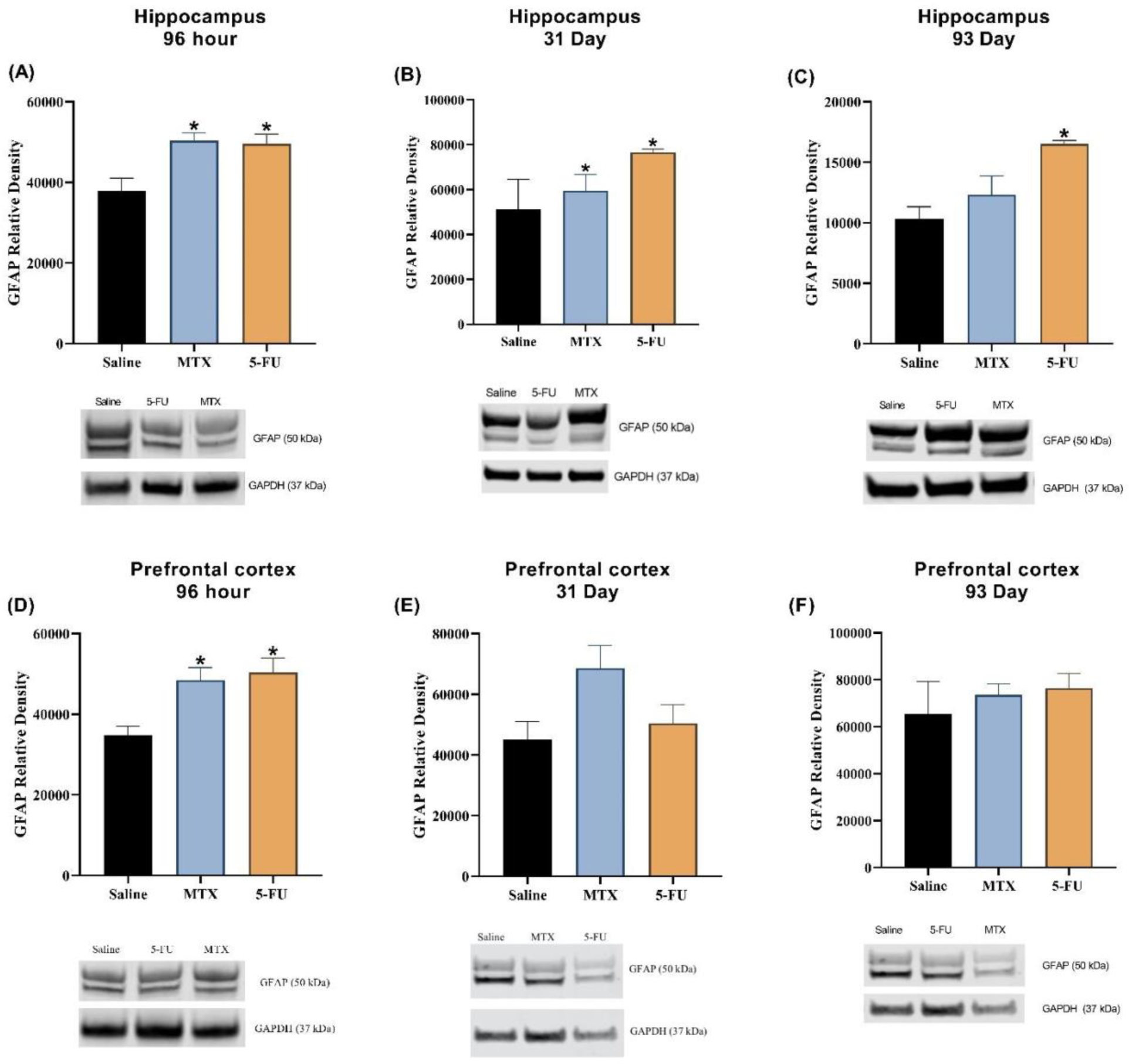
A-C: Western blot analysis of the expression of GFAP in the HIPP in Saline (n=5-6), 5-FU (n=5-6) and MTX (n=5-6) treated female SD rats at 96-hours (A), 31-days (B), and 93-days (C). Elevated GFAP was seen in animals treated with MTX and 5-FU at 96-hours and 31-days, but only in 5-FUanimals at 93-days. D-F: GFAP expression in the PFC in Saline (n=6), 5-FU (n=6) and MTX (n=6) treated female SD rats at 96-hours (D), 31-days (E), and 93-days (F). Elevated GFAP was seen in animals treated with MTX and 5-FU at 96-hours, but not at 31- or 93-days. Data expressed as mean ± SEM. Representative n = 6 per treatment group. However, outliers were removed if value > 2 SDs from the mean (n=1 MTX removed at 96-hours hippocampus and n=1 5-FU removed at 93-day hippocampus). * p<0.05 compared with saline.

For the PFC, there was a significant treatment effect in the expression of GFAP at 96-hours (F(2,15 = 7.927, p=0.004). Post-hoc analysis revealed a significant increase in the 5-FU treated group compared to saline controls (p=0.006) and in the MTX treated group compared to saline controls (p=0.015). No significant difference was observed between 5-FU and MTX treated animals (p=0.896) (Figure 6D). At 31-days, there was no significant treatment effect found in the PFC, although there was a trend towards statistical significance (F(2,15 = 3.565, p=0.054) (Figure 6E). Similarly, at 93-days, there was no significant treatment effect found in the PFC (F(2,15 = 0.384, p=0.688) (Figure 6F).

## 4. Discussion

The aim of this study was to investigate the time course of chemotherapy induced cognitive impairment and any potential concomitant changes in levels of astrocyte reactivity in a rat model of CICI at an acute (96-hours), sub-acute (31-days) and chronic time-point (93-days) following final chemotherapy treatment. The results of the present study suggest that the administration of chemotherapy impairs cognitive function at the sub-acute time-point, but that these impairments are ameliorated at a more chronic time-point. Furthermore, the results imply that impairment is not limited to a single cognitive domain, as demonstrated by impairment of both cognitive adaptability and memory in the Barnes maze task and task acquisition in the 5CSRTT. In addition to the behavioural results, chemotherapy administration was shown to cause astrocytic reactivity marker expression changes at all three time-points, particularly in the HIPP, suggesting that, although cognitive changes were not present at all time-points, there may be underlying neuroinflammatory changes that could drive later re-emergence of cognitive deficits. As CICI has been shown to affect patients up to 20-years post-final treatment, it is important to investigate even later chronic time-points, as there is currently limited evidence in the preclinical literature for this. Therefore, a more complete picture of the time course of CICI and its related neuroinflammatory changes in preclinical models is necessary.

At the acute time-point (96-hours), chemotherapy had no impact on recognition memory, as assessed through the NOR test, in spite of increased astrocytic reactivity in both the HIP and PFC. This implies that, whilst there was at least some degree of neuroimmune activation occurring at this early time point, the changes were not substantial enough to cause cognitive deficits, at least on the single cognitive test assessed. The previous literature on NOR performance in the acute period following chemotherapy administration is mixed. Similar to our findings, several studies have reported no detrimental effects on cognition in the NOR in the period up to 7-days following chemotherapy administration (41, 42). For example, one study found that male Long Evans rats treated with MTX (250mg/kg, injected intraperitoneally 10-times twice a week for 2-weeks, followed by weekly injections) showed no difference in novel object exploration compared to controls at either 3 or 7-days post-treatment (42). Similarly, another study found that female C57BL/J mice treated with 5-FU (3 times weekly, 100mg/kg) performed on par with controls at 1-week post-treatment (41). Conversely, others have demonstrated that various chemotherapy agents, including 5-FU, cause impairments in preference index in the NOR at timepoints ranging from 24-hours up to 4-weeks post-treatment (43, 44). It is possible that differences in behavioural testing protocol, dosage/length of time of chemotherapeutic agent administration and strain differences between studies may account for the differences observed (45). Given these discrepancies, the use of additional cognitive assessments, such as the Morris water maze (MWM), may be needed in order to fully elucidate changes to cognitive function that appear in the period immediately following chemotherapy exposure. It is important to note, however, that selecting a test for assessment of cognitive function in rodents at acute time-points is challenging (46, 47). Many such tests require an intense training period, often for several days or weeks, precluding them from use at early time-points. The NOR is advantageous in this respect because of its spontaneous nature, requiring no such pre-training. However, the conflicting results summarised above suggest some concerns over its reproducibility, and concerns have been raised over its sensitivity, with one study advising that it may be best suited for assessment of severe, rather than more subtle, cognitive impairment (48).

While no evidence of cognitive impairment was noted at this early time-point, an increase in levels of GFAP were noted within both the HIPP and PFC for both treatment groups compared to saline controls. It has been well evidenced in preclinical models of CICI that acute inflammation is present following chemotherapy administration (28). For example, an elevation in pro-inflammatory cytokines and chemokines (15), a decline in anti-inflammatory cytokines (49), and an increased expression of markers of microglial activation (44) and astrocytic reactivity (50) have all been noted between 0-7-days following chemotherapy administration (28). This is in line with our findings, which confirm that levels of astrocytic reactivity are increased at this early time-point, indicative of overall increases in neuroinflammation. While this does not appear to be significant enough to lead to impairments in cognition at this time-point, it is also possible that acute cognitive impairments induced by neuroinflammation are more subtle and that the NOR may not be sensitive enough to detect these changes (48). Conversely, it may be that it takes time for cognitive impairment to manifest following the induction of neuroinflammation by chemotherapy exposure.

In line with this, while cognitive changes were not present at the acute time-point, by 31-days following chemotherapy exposure, impairments were observed across multiple domains, including spatial memory, cognitive adaptability and complex task acquisition, as demonstrated across both the Barnes maze and 5CSRTT. Previously, assessments of CICI utilising the 5CSRTT have employed a study design in which C57BL/6J mice underwent the training phase (task acquisition), followed by cisplatin chemotherapy treatment, and then performed the probe phase. This schedule resulted in no observed deficit in learning and task acquisition (51, 52). A novel approach used in the current study was that animals were treated with chemotherapy *prior* to undergoing the training phase. In doing this, we showed an impairment in task acquisition in animals treated with 5-FU, specifically as stimulus duration decreased (and cognitive load increased), with effects noted for stimulus durations of 20-seconds or less. This finding is in line with the clinical literature, where patients with CICI often report deficits in learning and task acquisition when cognitive load is higher (53, 54).

While similar effects on task acquisition were not noted for the Barnes maze, this may be because the task to be acquired does not vary between training days and therefore there is no increase in cognitive demand for animals between training days.

Interestingly, no differences were noted on any of the measured outcomes on probe day in the 5CSRTT for the sub-acute cohort, with performance in attention, impulsivity, response inhibition and processing speed spared in both MTX and 5-FU treated animals. While this may initially appear surprising, clinical research has demonstrated that engaging in mentally stimulating tasks can improve symptoms of cognitive impairment, including executive function, through repetitive tasks with adaptive difficulty level (53). As the 5CSRTT requires a time intensive, repetitive training period with hierarchical difficulty, it is plausible that chemotherapy treated animals may have cognitively benefitted from engagement with the 5CSRTT. In line with this, similar beneficial effects were not noted on the Barnes maze, where task acquisition does not have the same mentally stimulating nature. Instead, on the Barnes maze, we observed that animals, irrespective of treatment group, performed similarly during the first probe trial, with a high degree of variability between individual animals, as a major component of what they have previously learnt shifted (i.e. the location of the escape box). Within the second trial, however, saline treated animals were able to improve their performance and quickly find the new escape box location, whilst those previously exposed to both chemotherapies took significantly longer, indicating that their cognitive adaptability was impaired (37, 55). Similarly, on the second probe trial, chemotherapy treated animals took significantly longer than controls to recall the original location of the old escape box, suggesting that they were also experiencing deficits in spatial memory (37, 55). This is consistent with the results of a study by Demby et al., where C57BL/6J mice receiving 5mg/kg DOX chemotherapy (twice over 2-weeks) performed worse in both the initial identification of the escape hole and the latency to get to the escape hole at 6-weeks post chemotherapy in the

Barnes maze (56). It is important to note that not all prior studies have been consistent. For example, in C57BL/6J mice receiving 2mg/kg of DOX (weekly for 4 weeks), a study found that there was no significant difference in the latency to reach the escape box or the distance travelled between treated and saline control animals (49). Overall, however, the findings in our study do align with the previous literature, where impairment following both MTX (doses ranging from 40mg/kg to 250mg/kg) and 5-FU (100mg/kg) treatment have been noted up to 5-weeks following chemotherapy treatment across hippocampal-based tasks, such as the MWM, NOR and open field test (18, 41, 42, 57). In previous studies utilising the MWM in particular, chemotherapy treated animals showed no difference in the learning/re-learning of the task compared to control animals but displayed significant impairments in spatial memory consolidation and recall, as evidenced by longer latency times to cross the platform location 16-21 days post-treatment (50, 57).

Of note, astrocytic reactivity was also present at the sub-acute time point, however only in the HIPP, and not the PFC. The HIPP appears to be particularly vulnerable to chemotherapy, with a previous systematic review of the literature in breast cancer patients suggesting that chemotherapy causes impaired neurogenesis, reduced volume and structural abnormalities in this region, leading to subsequent effects on hippocampal-related cognition (58). Therefore, it is plausible that this heightened vulnerability led to the increase in astrocyte reactivity noted in this region, compared to the PFC, contributing to the impairments seen on hippocampal-dependent tasks, such as recalling the location of the old escape box on probe day. While this is not sufficient to explain the full range of cognitive deficits observed in the current study, it has been shown that chemotherapy-induced hippocampal damage is associated with impaired functional connectivity to other brain regions, including between the HIPP and PFC, leading to impairments in general executive function, working memory and attention in breast cancer patients treated with tamoxifen (59). If a similar disruption of connectivity between the HIPP and PFC occurs in rodents exposed to chemotherapy, it may help explain the impairments in cognitive adaptability observed in the current study. To confirm this, further research is warranted in both cancer survivors and animal models to explore alterations in connectivity between the HIPP and prefrontal circuits using neuroimaging techniques, such as diffusion tensor imaging or functional magnetic resonance imaging, in order to probe their potential contribution to cognitive impairments observed in CICI.

Given the impairments in cognitive function seen at the sub-acute time-point, we were surprised that, at 93-days following exposure, there was no impact of chemotherapy on any aspect of cognitive performance assessed, with the exception of an increase in correct response latency at only the 2.5s stimulus duration in 5-FU treated animals compared to saline controls. This suggests that CICI is largely resolved by this chronic time-point. To date, assessment of chronic time-points has been less performed in preclinical CICI studies, making comparisons and conclusions between our results and those of previous studies difficult. Nevertheless, our findings are consistent with those of several previous preclinical studies. Seigers et al. 2015 demonstrated that male C57BL/J mice treated with commonly used chemotherapy agents docetaxel, doxorubicin, cyclophosphamide, 5-FU, topotecan or MTX displayed short-term cognitive impairments in tests such as the open field test, NOR and Barnes maze, up to 2-weeks post-treatment, but did not have any long-term detrimental effects at 15-weeks post-treatment (60). Similarly, Borbélyová et al. 2020 reported cognitive impairment in the T-maze immediately after treatment with multi-agent chemotherapy, which was not present at 3-months post-treatment (61). Whilst these findings are similar to those observed in this study, they do not align well with the clinical literature, where cognitive impairment is evident up to 20-years post-treatment (62). Differences could be attributed to the fact that many CICI animal models are tumour free and therefore do not account for how cancer itself may contribute to cognitive impairments. Additionally, treatment regimens differ widely in terms of dose and type of chemotherapy administered; therefore, further work is required to probe these chronic time-points in animal models of CICI, in order to establish a gold standard animal model, which is more reflective of the human condition.

Importantly, whilst there was no evidence of cognitive impairment at this time-point, levels of astrocytic reactivity were still significantly increased in the HIPP, but not the PFC, of animals treated with 5-FU. This could suggest that some degree of neuroinflammation is still persistent at this time-point, at least in certain regions of the brain, and could contribute to the re-emergence of cognitive impairment at later time-points. Conversely, elevations in GFAP were not noted in either brain region in animals treated with MTX. This may be due to a difference in the mechanism of action of the chemotherapeutic agents administered. In contrast to MTX, 5-FU can penetrate the BBB (63, 64) and therefore has the potential to cause direct insult to the CNS, even when administered systemically (43, 65). As previously described, the HIPP is particularly susceptible to chemotherapy-induced damage and this, in conjunction with the notable neurotoxic potential of 5-FU, may account for why inflammation is still persistent at the chronic time-point in this region in 5-FU-treated animals (43, 65). This may also help to explain why, in general, more significant impairment was elucidated by 5-FU than by MTX.

One limitation of the current work is that the assessment of neuroinflammation was limited to protein expression levels of GFAP. It is important to note that the expression pattern of neuroimmune markers overall is not well characterised in CICI. Particularly, the level of expression change that is defined as relevant, or exceeding the ‘normal’ threshold, needed to manifest into pathological effect, such as cognitive impairment, has not been clearly established in this context (28, 66, 67). Additionally, although GFAP is a commonly utilised marker of astrocytic reactivity and is useful to assess in animal models of injury and disease, when assessed alone, it is only a fraction of the complex neuroinflammatory process that is occurring (68). Markers of microglial activation (e.g. CD-68) or phenotype (e.g. MHCII vs MHCI) would also prove useful, in order to provide context on the pro- or anti-inflammatory nature of neuroimmune activation following chemotherapy exposure, as not all inflammation is detrimental. Furthermore, assessing a greater array of neuroimmune markers would allow for a more comprehensive insight into if, when, and where neuroinflammation may be occurring in CICI. This is a critical first step in treatment development.

The animal model itself also represents a potential limitation of the current work. Currently, there is no gold standard model of CICI utilised within the field, with several different strains, types of chemotherapies, doses administered and behavioural paradigms utilised and various time-points investigated. This makes it difficult to compare between the results of individual studies. It is possible that, given the relatively subtle cognitive deficits observed, particularly at the chronic time-point, the model used in this study did not elucidate CICI to a great enough extent to be comparable to that seen clinically, and therefore a model that elicits more severe cognitive impairment may be needed. Further work needs to be conducted in order devise a model where the optimal dose of chemotherapy is determined, potentially through conducting a dose response study in order to elucidate when specifically cognitive changes begin to emerge following chemotherapy and to characterise the nature and severity of such changes. This study also focuses solely on young female rats, limiting its scope to a specific age and sex. Future research should encompass a broader range of ages and include both sexes to better understand the clinical implications and whether there are sex and age differences.

## 5. Conclusion

This study has identified that treatment with chemotherapeutic agents MTX and 5-FU impairs cognitive function at a sub-acute time-point following treatment, with impairment not limited to a single cognitive domain. Interestingly, this impairment was ameliorated at the chronic time-point, suggesting either transient effects or a potential activation of a repair mechanism. In contrast, this study demonstrated that levels of astrocytic reactivity occurred at all time-points studied, particularly within the HIPP, suggesting that the presence of neuroinflammation does not always indicate that cognitive impairment will occur in tandem. This highlights the importance of investigating other neural mechanisms that might contribute to the emergence or maintenance of CICI. Our findings provide valuable insight into the complexity of the neuroinflammatory response in CICI, and its implications for cognitive function. They also highlight the need for further investigation to probe the complexity of neuroinflammation in CICI. Such work is a critical first step in understanding which specific aspects of this response to target and at what time-points, which is significant, as neuroinflammation may be a promising target to treat CICI and increase quality of life for cancer survivors.

## Acknowledgements and Funding

This study was funded by the Neurosurgical Research Foundation. ALW was supported by a NHMRC Peter Doherty Biomedical Research Fellowship (APP1140072). OJH, IS and RPG were supported by an Australian Government Research Training Program Scholarship.

## 7. Ethics Statement

All experimental procedures were approved by the University of Adelaide Animal Ethics Committee (S-2019-019) and were in accordance with the Australian Code for the Care and Use of Animals for Scientific Purposes, 8^th^ Edition, 2013 guidelines.

## 8. Author Contributions

RPG, IS and OJH contributed to the conception, design, data acquisition, analysis, interpretation and manuscript preparation. ALW and LCP contributed to the conception, design, analysis, interpretation and manuscript preparation. All authors contributed to the manuscript and approved the final submission.

